## Supplemental Data for "External location of touch is constructed post-hoc based on limb choice"

### Supplementary Information

#### Experiment 1

Supplementary Table 1. Generalized Linear Mixed Models of temporal order judgment accuracy in Experiment 1.

| | df | $\chi^2$ | p |
|---|---|---|---|
| all trial phases (full model) | | | |
| start posture | 16, 17 | 22.03 | <0.001 |
| end posture | 16, 17 | 83.82 | <0.001 |
| movement phase | 14, 17 | 60.90 | <0.001 |
| start posture : end posture | 16, 17 | 9.91 | <0.01 |
| start posture : movement phase | 14, 17 | 9.72 | .02 |
| end posture : movement phase | 14, 17 | 13.86 | <0.01 |
| start posture : end posture : movement phase | 14, 17 | 31.59 | <0.001 |
| phase 1: before movement onset | | | |
| start posture | 4, 5 | 23.09 | <0.001 |
| end posture | 4, 5 | 8.48 | 0.004 |
| start posture : end posture: | 4, 5 | < 0.01 | 0.96 |
| phase 2: first half of movement | | | |
| start posture | 4, 5 | 11.72 | <0.001 |
| end posture | 4, 5 | 30.61 | <0.001 |
| start posture : end posture: | 4, 5 | 0.11 | 0.74 |
| phase 3: second half of movement | | | |
| start posture | 4, 5 | 0.69 | 0.41 |
| end posture | 4, 5 | 55.59 | <0.001 |
| start posture : end posture: | 4, 5 | 44.22 | <0.001 |
| phase 4: after movement offset | | | |
| start posture | 4, 5 | 4.15 | 0.04 |
| end posture | 4, 5 | 22.67 | <0.001 |
| start posture : end posture: | 4, 5 | 2.33 | 0.13 |

##### Movement times

A LMM with the factor Posture (uncrossed-uncrossed, uncrossed-crossed, crossed-uncrossed, crossed-crossed) on movement times revealed a significant main effect ($\chi^2(3,6) = 38.34$, $p < 0.001$). Post hoc test (Bonferroni corrected) showed that movement times were significantly longer in the crossed-uncrossed condition compared to the other three conditions (all $p < 0.001$).

Supplementary Table 2. Movement times (in ms) in Experiment 1

| Posture | Mean | 95% CI Range |
|---|---|---|
| Uncrossed-uncrossed | 500 | 453 – 548 |
| Uncrossed-crossed | 487 | 440 – 535 |

| | | |
|---|---|---|
| Crossed-uncrossed | 558 | 510 – 605 |
| Crossed-crossed | 475 | 427 – 522 |

### Experiment 2

#### Movement times

A LMM with the factors Posture (uncrossed-uncrossed, crossed-crossed) and SOA (60 ms, 85 ms, 110 ms, 135 ms) showed that movement times were statistically similar across conditions – main effect of posture: $\chi^2(9,10) = 1.31$, $p = 0.25$; main effect of SOA: $\chi^2(7,10) = 0.03$, $p = 0.99$; interaction: $\chi^2(7,10) = 0.09$, $p = 0.99$.

Supplementary Table 3. Movement times (in ms) in Experiment 2

| Posture | SOA | Mean | 95% CI Range |
|---|---|---|---|
| Uncrossed-uncrossed | 60 ms | 563 | 523 – 604 |
| Uncrossed-uncrossed | 85 ms | 562 | 522 – 603 |
| Uncrossed-uncrossed | 110 ms | 563 | 522 – 604 |
| Uncrossed-uncrossed | 135 ms | 563 | 522– 603 |
| Crossed-crossed | 60 ms | 571 | 531 – 612 |
| Crossed-crossed | 85 ms | 570 | 530 – 611 |
| Crossed-crossed | 110 ms | 569 | 528 – 609 |
| Crossed-crossed | 135 ms | 571 | 530 – 612 |

#### Hand Assignment

TOJ performance in Experiment 2 was modulated by hand posture and SOA (see Supplementary Figure 1). A GLMM with factors Posture (uncrossed-uncrossed, crossed-crossed) and SOA (60 ms, 85 ms, 110 ms, 135 ms) revealed significant main effects of Posture ($\chi^2(8,9) = 586.94$, $p < 0.001$) and SOA ($\chi^2(6,9) = 218.00$, $p < 0.001$), and a significant interaction ($\chi^2(6,9) = 66.63$, $p < 0.001$). Post hoc analysis of the interaction (Bonferroni corrected, Supplementary Table 2) showed that TOJ performance was better when the arms were in an uncrossed compared to a crossed posture at all SOAs. Furthermore, performance increased with SOA duration for the uncrossed posture but was relatively similar across all SOAs for the crossed posture.

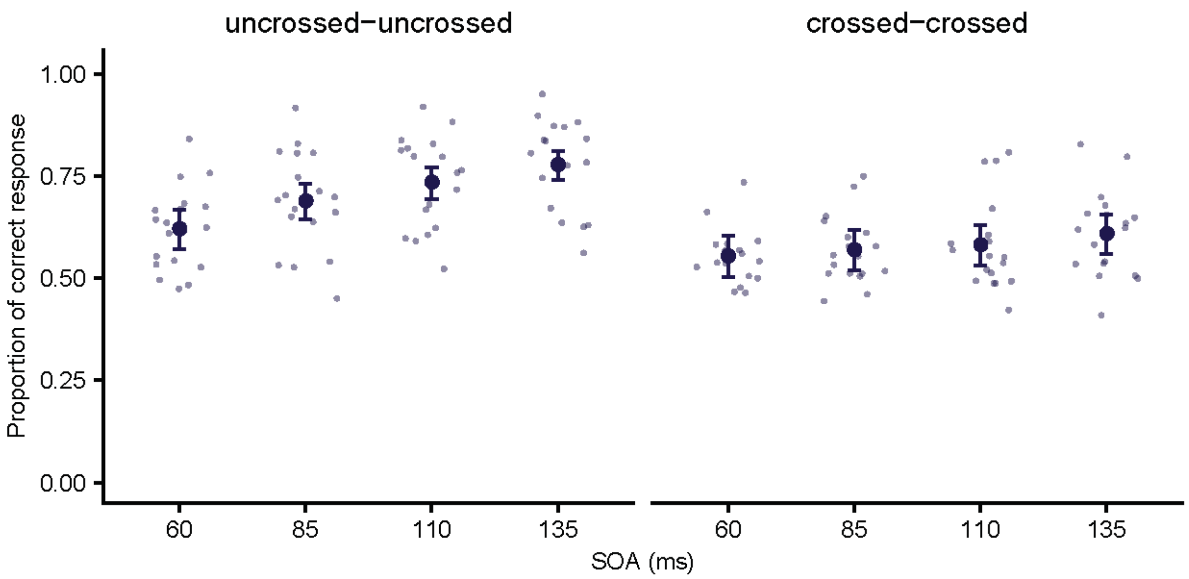

**Supplementary Figure 1.** Proportion of correct hand assignment across movement conditions (uncrossed-uncrossed, crossed-crossed) and SOA (60ms, 85ms, 110ms, 135ms) in Experiment 2. Error bars denote 2 s.e. from the mean; asymmetry is due to nonlinear conversion from the GLMM's logit scale to percentage correct. Large symbols are group means, small symbols are individual participants' performance.

Supplementary Table 4. Post hoc analysis of temporal order judgment accuracy in Experiment 2.

| **Contrast** | **Estimate** | **SE** | **df** | **Z** | **P** |
|---|---|---|---|---|---|
| uu,60 - cc,60 | 0.3237 | 0.0509 | Inf | 6.356 | <.0001 |
| uu,60 - uu,85 | -0.3398 | 0.0518 | Inf | -6.563 | <.0001 |
| uu,60 - uu,110 | -0.5715 | 0.0532 | Inf | -10.746 | <.0001 |
| uu,60 - uu,135 | -0.8192 | 0.0550 | Inf | -14.886 | <.0001 |
| cc,60 - cc,85 | -0.0867 | 0.0515 | Inf | -1.684 | 1.0000 |
| cc,60 - cc,110 | -0.1078 | 0.0515 | Inf | -2.095 | 1.0000 |
| cc,60 - cc,135 | -0.2587 | 0.0518 | Inf | -4.991 | <.0001 |
| uu,85 - cc,85 | 0.5768 | 0.0524 | Inf | 11.003 | <.0001 |
| uu,85 - uu,110 | -0.2317 | 0.0546 | Inf | -4.245 | 0.0006 |
| uu,85 - uu,135 | -0.4794 | 0.0564 | Inf | -8.502 | <.0001 |
| cc,85 - cc,110 | -0.0212 | 0.0515 | Inf | -0.411 | 1.0000 |
| cc,85 - cc,135 | -0.1720 | 0.0519 | Inf | -3.317 | 0.0255 |
| uu,110 - cc,110 | 0.7873 | 0.0538 | Inf | 14.631 | <.0001 |
| uu,110 - uu,135 | -0.2476 | 0.0577 | Inf | -4.293 | 0.0005 |
| cc,110 - cc,135 | -0.1508 | .05190 | Inf | -2.908 | 0.1018 |
| uu,135 - cc,135 | 0.8841 | 0.0560 | Inf | 15.799 | <.0001 |

#### Explicit stimulus localization in space

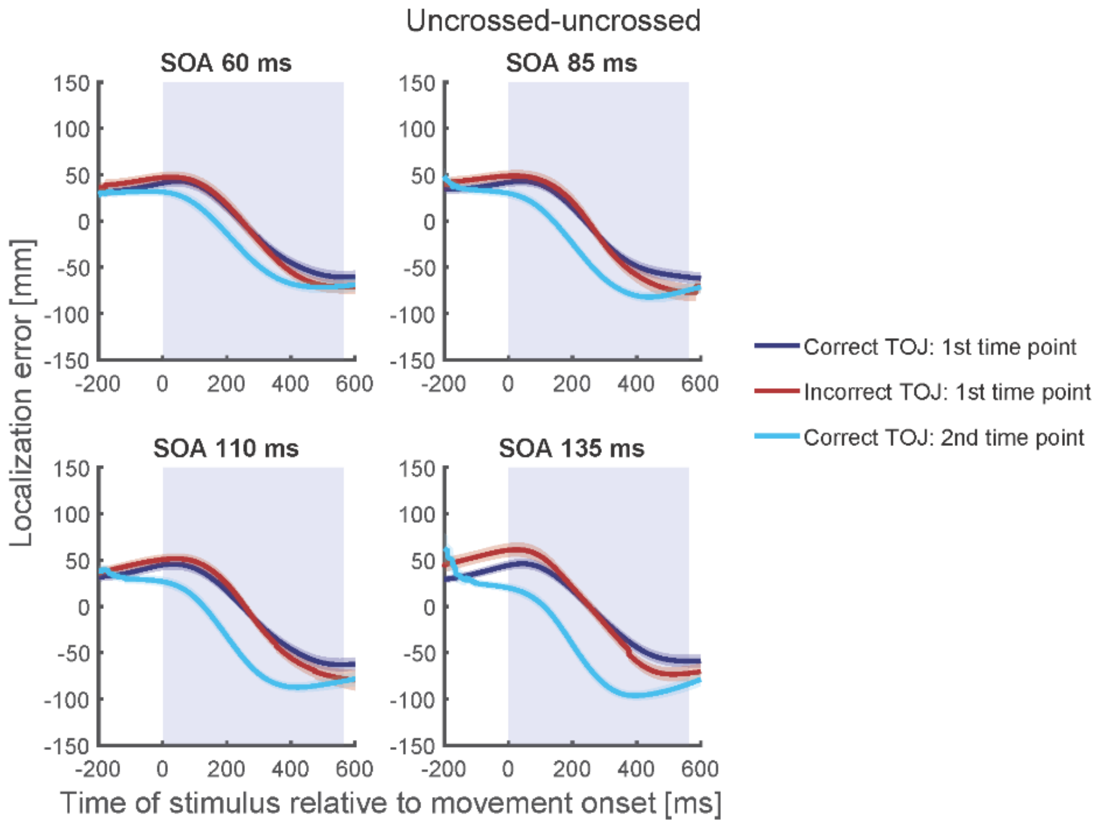

***Supplementary Figure 2.*** *Localization curves of the uncrossed-uncrossed posture condition, averaged across participants, for each of the four SOAs in Experiment 2. Curves of incorrect TOJ trials (red) show a similar pattern as the localization curves of the correct TOJ trials at time 1 (dark blue), but not as the localization curves of the correct TOJ trials at time 2 (light blue). Traces reflect the mean, shaded areas around the traces reflect s.e.m. The shaded regions in the background represent the average movement time.*

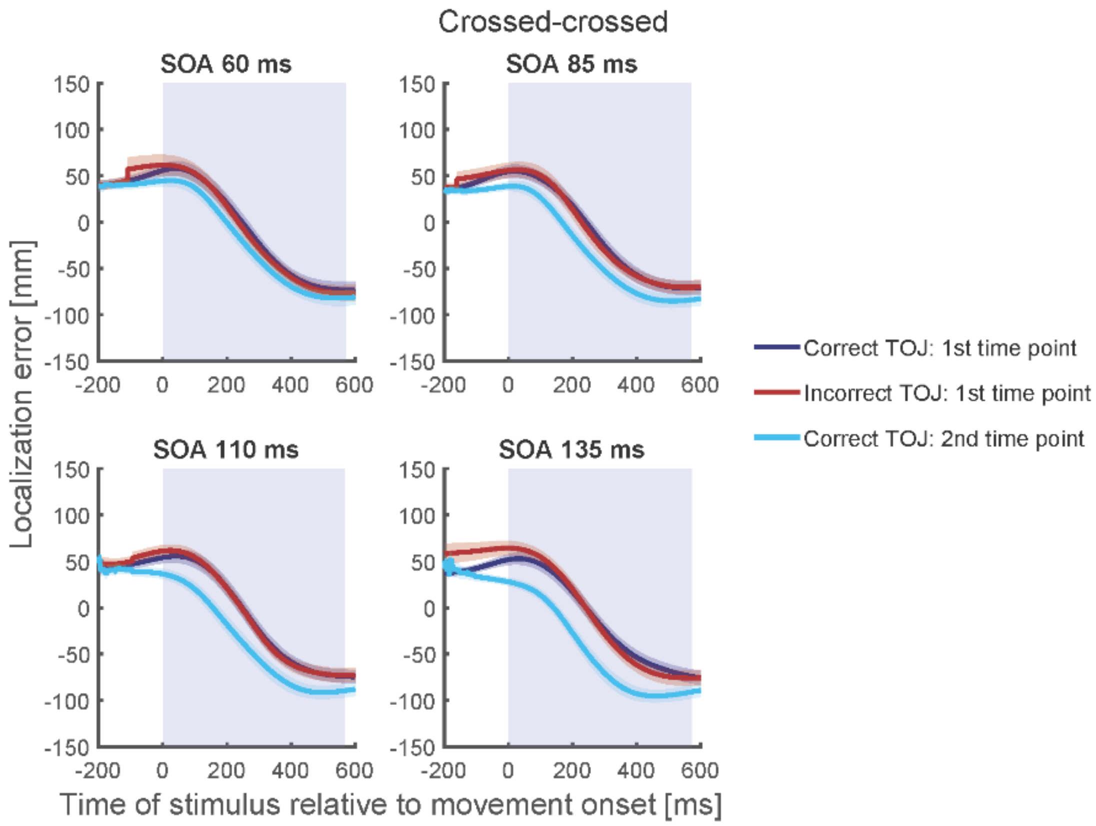

***Supplementary Figure 3.*** *Localization curves of the crossed-crossed posture condition, averaged across participants, for each of the four SOAs in Experiment 2. Curves of incorrect TOJ trials (red) show a similar pattern as the localization curves of the correct TOJ trials at time 1 (dark blue), but not as the localization curves of the correct TOJ trials at time 2 (light blue). Traces reflect the mean, shaded areas around the traces reflect s.e.m. The shaded regions in the background represent the average movement time.*

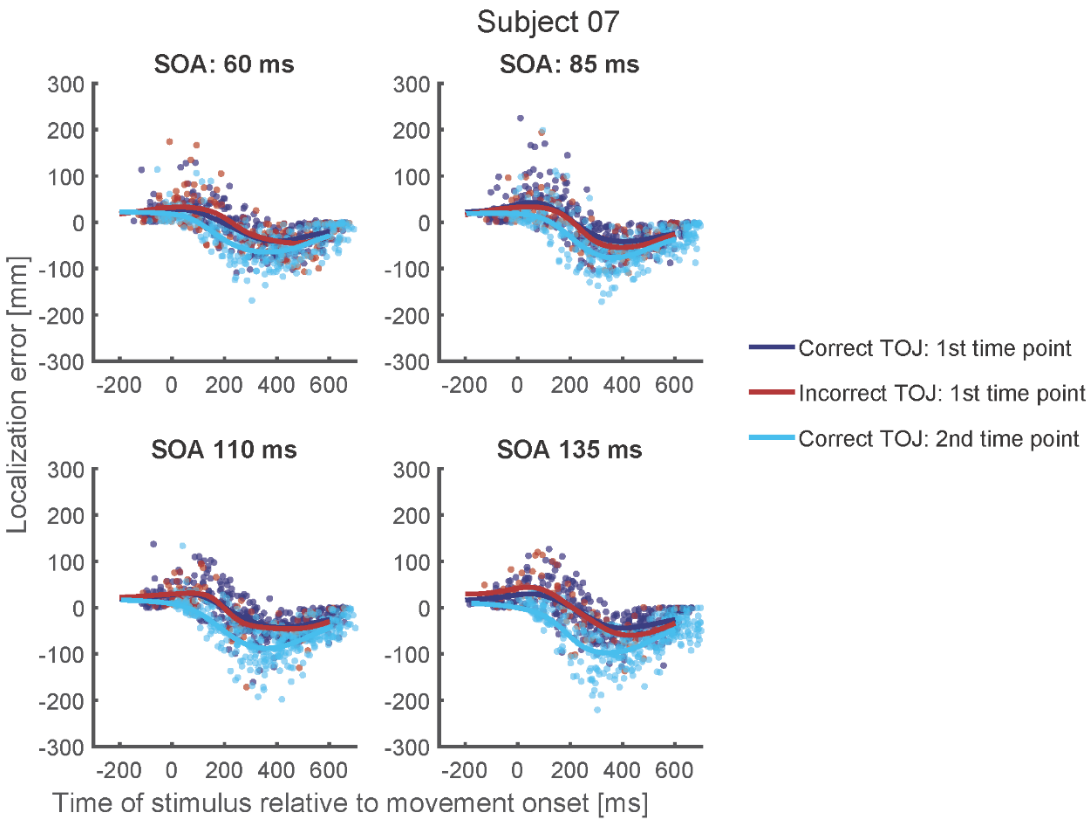

***Supplementary Figure 4.*** *Localization curves of a representative participant (#07), averaged across posture, for each of the four SOAs in Experiment 2. Curves of incorrect TOJ trials (red) show a similar pattern as the localization curves of the correct TOJ trials at time 1 (dark blue), but not as the localization curves of the correct TOJ trials at time 2 (light blue).*

Supplementary Table 5. Bayesian model estimates for Experiment 2

| **Time 1 (ms)** | | | |
|---|---|---|---|
| **Model** | **Intercept** | **Error** | **95% CI Range** |
| Common Intercept: [shift ~ 1 + (1 I participant )] | 7.79 | 7.44 | -6.77 – 22.66 |
| Individual Intercept per SOA: [shift ~ SOA + (1 I participant )] | | | |
| SOA 60 ms | 10.83 | 9.98 | -8.50 – 30.65 |
| SOA 85 ms | 4.33 | 9.88 | -14.95 – 24.11 |
| SOA 110 ms | 0.32 | 10.20 | -19.16 – 20.52 |
| SOA 135 ms | 15.54 | 10.14 | -4.31 – 35.53 |
| **Time 2 (ms)** | | | |
| **Model** | | | |
| Common Intercept: [shift ~ 1 + (1 I participant )] | -78.84 | 9.92 | -98.23 – -58.64 |
| Individual Intercept per SOA: [shift ~ SOA + (1 I participant )] | | | |
| SOA 60 ms | -51.43 | 13.33 | -77.93 – -25.38 |
| SOA 85 ms | -74.83 | 13.42 | -102.03 – -48.78 |
| SOA 110 ms | -87.33 | 13.49 | -114.71 – -61.85 |
| SOA 135 ms | -104.56 | 13.39 | -131.39 – -79.12 |
